# DynoSys 2.0: Graph-Based Modeling of Dynamic Risk States and System Transitions in Human Behaviours Development

**DOI:** 10.64898/2026.05.06.723259

**Authors:** Mengman Wei, Qian Peng

## Abstract

Human behavioral and mental health outcomes arise from interactions among genetic, environmental, and neurobiological systems. Existing frameworks often model these components jointly, but many treat variables independently or use static representations. This limits their ability to capture system-level dynamics and changes over time. To address this, we developed DynoSys, a unified framework that integrates these signals using three layers: predictive models, relationship exploration models, and mechanism-oriented explanation models. Building on this framework, we introduce DynoSys 2.0, a graph-based temporal modeling approach inspired by the free-energy principle by Karl Friston. In this framework, each individual is represented as a dynamic graph that evolves over time. We hypothesize that healthy development and adverse mental health outcomes correspond to different system states and trajectories.

Using longitudinal data from the Adolescent Brain Cognitive Development (ABCD) Study, we construct time-indexed graphs that integrate polygenic risk scores (PRS), multi-domain environmental features, and neuroimaging-derived representations. We study six phenotypes: externalizing behavior, internalizing behavior, and substance use initiation (alcohol, nicotine, cannabis, and any substance). In these graphs, nodes represent domain-level features, and edges capture relationships derived from data-driven feature selection and temporal dependencies. We model graph evolution using recurrent neural networks and graph-temporal learning methods. We also define system-level measures, including graph energy and state transitions, to quantify dynamic patterns. Our results show that DynoSys 2.0 can model behavioral development using longitudinal multi-domain data. The framework achieved meaningful prediction for both continuous behavioral symptoms and substance-use initiation outcomes, but performance differed by outcome type. Externalizing behavior was predicted more accurately than internalizing behavior, and alcohol and any substance initiation showed stronger prediction than cannabis and nicotine initiation. Graph-derived energy measures showed clearer separation for high-versus low-symptom externalizing and internalizing groups, suggesting that continuous behavioral symptoms may be linked to different latent system states over time. Overall, DynoSys 2.0 provides a flexible framework for studying behavioral risk as a dynamic developmental process, while rare-event prediction and detailed graph-level interpretation require further work.

## 1 Introduction

Human development can be viewed as a dynamic system shaped by interactions among biological, environmental, and neurobehavioral domains. In our previous work, the DynoSys framework [1] was developed to model this system using longitudinal and multimodal data. This framework integrates genetic, environmental, and neurobiological factors to predict behavioral outcomes.

However, the original DynoSys framework represents variables mainly as independent features or aggregated domain-level summaries. Although this approach is effective for prediction, it does not explicitly capture relationships between variables or how these relationships change over time.

Many complex systems are more naturally represented as networks or graphs, where nodes represent system components and edges represent interactions [2]. In neuroscience, network-based approaches have been widely used to study brain connectivity [3]. In systems biology, graph models are used to represent interactions among genes and proteins [2]. These studies show that graph-based representations are useful for understanding complex, interconnected systems.

Extending this idea to human development, individuals can be represented as dynamic graphs in which multiple domains interact and evolve over time. This allows both relationships and temporal changes to be modeled in a unified way.

In addition, theoretical work suggests that healthy and unhealthy conditions may correspond to different system states or trajectories. The free-energy principle proposes that biological systems maintain stability by minimizing internal uncertainty, and deviations from this process may reflect abnormal system states [4]. This perspective highlights the importance of studying system-level behavior rather than focusing only on individual predictors.

To address these limitations, we propose DynoSys 2.0, a graph-based extension of the original framework. As shown in Figure 1, this approach introduces three main components:

1. **Graph representation of multimodal systems**. Variables are organized as nodes and edges to capture relationships across domains.
2. **Temporal modeling of graph evolution**. Graph structures change over time, allowing the study of developmental trajectories.
3. **System-level state analysis**. Graph-based metrics are used to measure system stability, transitions, and risk states.

**Figure 1.**
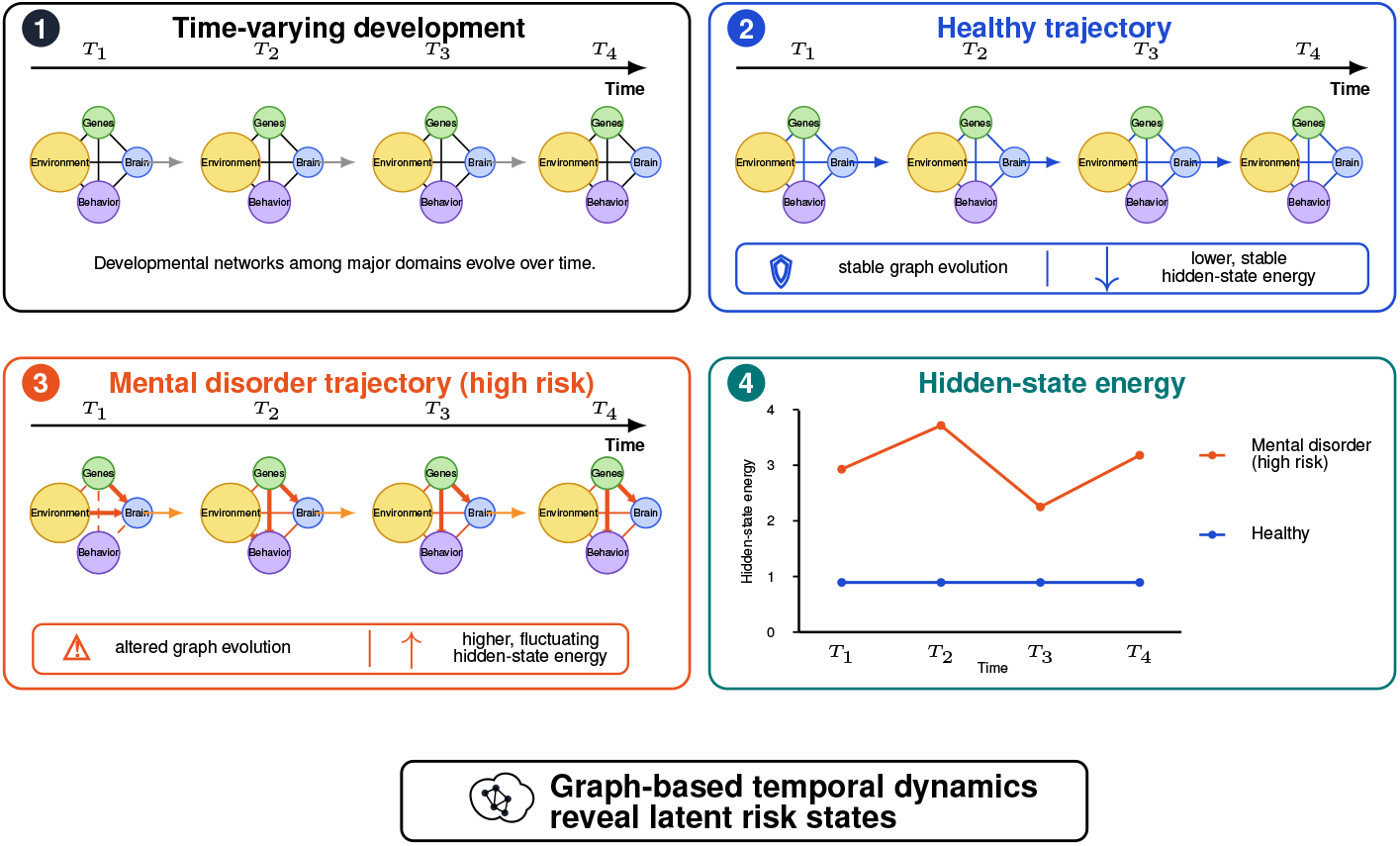
Conceptual framework of graph-based temporal development and latent system energy in DynoSys 2.0. Each individual is represented as a time-varying graph across developmental time points *T*_1_, *T*_2_, *T*_3_, and *T*_4_. Nodes represent major developmental domains, including genes, environment, brain, and behavior. Edges represent relationships among these domains and are allowed to change over time. In the healthy trajectory, graph structure remains relatively stable across development and is associated with lower, more stable hidden-state energy. In the mental disorder or high-risk trajectory, graph structure changes more irregularly, with stronger or dysregulated connections and higher, more fluctuating hidden-state energy. The hidden-state energy plot summarizes these latent system differences over time. Overall, the framework proposes that graph-based temporal dynamics can reveal latent risk states that distinguish healthy development from adverse mental health trajectories.

Together, this framework moves beyond static prediction and enables the study of dynamic system behavior over time.

## 2 Methods

### 2.1 Study Design and Overall Framework

This study extends the DynoSys framework to a graph-based temporal modeling setting. The goal is to model human development as a dynamic system integrating genetic, environmental, neurobiological, and behavioral data over time. Each individual is represented as a longitudinal sequence of system states, and both feature-level and graph-level temporal dynamics are modeled.

The overall design follows principles from systems neuroscience and multimodal integration frameworks, where complex behaviors arise from interactions across multiple domains rather than isolated predictors [5, 6].

The full pipeline consists of four stages:

1. unified longitudinal data construction;
2. sequence and graph representation building;
3. temporal modeling using LSTM and graph-temporal models;
4. system-level state analysis.

All steps were implemented in a reproducible pipeline using high-performance computing.

### 2.2 Data Sources and Phenotypes

Data were derived from the Adolescent Brain Cognitive Development (ABCD) Study, a large-scale longitudinal study designed to characterize brain development and child health [7, 8, 9].

The following modalities were included:

- **Genetic data:** polygenic risk scores (PRS);
- **Environmental data:** multi-domain longitudinal exposures;
- **Neuroimaging data:** brain-derived features;
- **Behavioral outcomes:** continuous and event-based developmental phenotypes.

Two types of outcomes were modeled. Continuous outcomes included externalizing behavior and internalizing behavior. Binary time-to-event outcomes included alcohol initiation, nicotine initiation, cannabis initiation, and any substance-use initiation.

### 2.3 Longitudinal Panel Construction

#### 2.3.1 Time Indexing

All data were aligned across discrete developmental time points: baseline, 1-year follow-up, 2-year follow-up, 3-year follow-up, and 4-year follow-up. These were mapped to integer time indices:

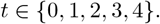

Each subject contributes repeated observations over time, consistent with longitudinal panel modeling approaches [10].

#### 2.3.2 Continuous Outcome Construction

For continuous phenotypes, the observed score *y*_*i,t*_ was directly used. Temporal features were derived as follows:

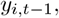

and

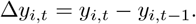

The lagged outcome and change score capture temporal dependencies and developmental trajectories [11].

#### 2.3.3 Binary Outcome Construction

For substance initiation outcomes, a discrete-time survival formulation was used [12]. Let *T*_*i*_ denote the time to event and *δ*_*i*_ denote the event indicator. The longitudinal binary outcome was defined as:

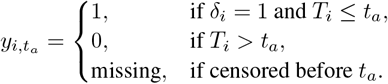

This produces a discrete-time hazard representation aligned with longitudinal covariates.

### 2.4 Feature Construction

#### 2.4.1 Polygenic Risk Score Construction

Polygenic risk scores were constructed to capture genetic liability across three major domains: externalizing behavior, internalizing psychopathology, and substance use.

For the externalizing domain, PRS were derived from large-scale multivariate genome-wide association studies of externalizing behaviors [13, 14]. For the internalizing domain, multiple PRS were constructed based on GWAS of major depressive disorder, anxiety disorders, neuroticism, and youth internalizing symptoms [15, 16, 17, 18]. To improve robustness and better reflect the multidimensional nature of internalizing traits in a youth sample, a composite internalizing PRS was additionally generated by integrating these related phenotypes.

For the substance-use domain, PRS were constructed using GWAS summary statistics for problematic alcohol use, cannabis use disorder, nicotine and alcohol use, and general substance-use disorder liability [19, 20, 21, 22]. For each GWAS dataset, summary statistics were harmonized and reformatted for compatibility with PRS-CS [23]. Posterior single nucleotide polymorphism effect sizes were estimated separately for each chromosome using linkage disequilibrium information from an external reference panel and then combined to generate genome-wide effect-size estimates.

Individual-level PRS were calculated using PLINK 1.9 [24] by summing allele dosages weighted by PRS-CS-derived effect sizes. All PRS were computed at the individual level, aligned to a consistent genotype reference panel, and standardized prior to downstream analyses when appropriate.

#### 2.4.2 Covariates

Covariates included sex, age, genetic principal components PC1–PC10, and site indicators. Adjustment using genetic principal components accounts for population stratification in genetic analyses [25].

#### 2.4.3 Domain-Level Features

Features were organized into predefined domains, including environment, family context, physical health, technology use, neurocognition, and brain. These domains were designed to capture distinct aspects of the developmental system, covering both external exposures and internal functional characteristics. The detailed construction and variable definitions for each domain follow the procedures described in the original DynoSys framework [1], ensuring consistency and comparability across studies.

#### 2.4.4 Brain Features

Neuroimaging features were incorporated at multiple developmental time points, including baseline, 24-month follow-up, and 48-month follow-up. This allowed the modeling of both static and longitudinal brain characteristics.

To obtain compact and interpretable representations, multiple types of feature summaries were derived from the imaging data, including principal component analysis-based representations, weighted feature scores, and cluster-based features. Principal component analysis was applied as a dimensionality reduction technique to capture major sources of variation in high-dimensional neuroimaging data, a widely adopted approach in neuroimaging studies [26].

### 2.5 Data Preprocessing

#### 2.5.1 Missing Value Handling

Missing values were imputed using median imputation:

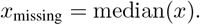

This approach is robust to outliers and commonly used in high-dimensional biomedical data [27].

#### 2.5.2 Standardization

All features were standardized:

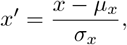

where *µ*_*x*_ and *σ*_*x*_ denote the mean and standard deviation of feature *x*, respectively.

#### 2.5.3 Residualization Against Covariates

Predictors were residualized to remove confounding effects:

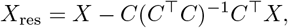

where *X* denotes the predictor matrix and *C* denotes the covariate matrix. This corresponds to linear regression-based adjustment [28].

### 2.6 Sequence Representation

Each subject was represented as a temporal sequence:

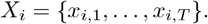

Sequences were padded and masked:

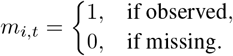

This representation is standard in sequence modeling [29].

### 2.7 Graph Construction

#### 2.7.1 Node Definition

Nodes represent domain-level feature groups, including genetic, environmental, neurobiological, and behavioral domains.

#### 2.7.2 Edge Construction

Edges were represented using an adjacency matrix:

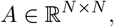

where *N* denotes the number of nodes. Graph representations follow network modeling approaches in neuroscience and systems biology [5].

### 2.8 Temporal Modeling

#### 2.8.1 LSTM Model

Recurrent neural networks were used to model temporal dependencies using Long Short-Term Memory units [30]. The hidden state update was defined as:

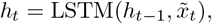

and the prediction was defined as:

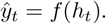

For continuous outcomes, the model was trained using mean squared error loss. For binary outcomes, the model was trained using cross-entropy loss.

#### 2.8.2 Graph-Temporal Model

Graph structure was integrated through message passing:

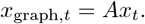

Temporal dynamics were then modeled using gated recurrent units [31]. This combines graph learning and temporal modeling [32].

### 2.9 Model Training

Models were trained using the Adam optimizer [33]. Early stopping was applied based on validation performance to reduce overfitting [34].

### 2.10 Evaluation Metrics

For continuous outcomes, model performance was evaluated using root mean squared error, mean absolute error, coefficient of determination, and Pearson correlation:

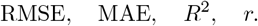

For binary outcomes, model performance was evaluated using area under the receiver operating characteristic curve, area under the precision-recall curve, and Brier score:

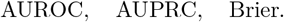

These are standard metrics for regression and classification tasks [35].

### 2.11 Graph State Analysis

Graph energy was defined as:

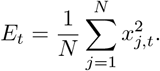

Graph-propagated energy was defined as:

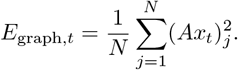

Temporal change in energy was defined as:

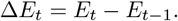

A collapse-like state was defined as:

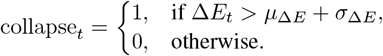

This definition is inspired by dynamical systems and stability analysis [36].

## 3 Results

### 3.1 Longitudinal sequence construction and model inputs

We constructed longitudinal sequence inputs for six behavioral outcomes, including two continuous symptom trajectories, externalizing and internalizing behavior, and four substance-use initiation outcomes: alcohol, nicotine, cannabis, and any substance initiation. Across all five random seeds, the dataset was split into 1,676 training subjects, 420 validation subjects, and 524–525 test subjects, depending on the outcome. All phenotypes were represented using 11 system-level nodes and up to five longitudinal time points, providing a consistent dynamic input structure across outcomes.

The number of longitudinal rows differed by outcome type. Continuous behavioral outcomes contained more repeated observations, with approximately 7,088 training rows and 2,227 test rows for externalizing and internalizing behavior. In contrast, substance-use initiation outcomes contained 5,028 training rows and 1,572 test rows, reflecting the event-based structure of the initiation phenotypes. Feature dimensionality also varied across outcomes, ranging from 39 features for nicotine initiation to 145 features for any substance initiation. These results indicate that DynoSys 2.0 generated stable and reproducible longitudinal sequence inputs across seeds while preserving outcome-specific differences in feature availability and target structure.

### 3.2 Predictive performance across continuous behavioral outcomes

For continuous behavioral trajectories, Graph-LSTM outperformed Graph-GTRNN for both externalizing and internalizing behavior (Table 2). Externalizing behavior was predicted with the highest accuracy among the continuous outcomes. Graph-LSTM achieved a mean *R*^2^ of 0.279 ± 0.017, RMSE of 8.237 ± 0.080, and Pearson correlation of 0.532 ± 0.015 across five seeds. In contrast, Graph-GTRNN showed weaker performance for externalizing behavior, with a mean *R*^2^ of 0.066 ± 0.099, RMSE of 9.368 ± 0.495, and correlation of 0.129 ± 0.268.

**Table 1.** Cohort, outcome structure, and model input summary across six phenotypes.

| Outcome | Prediction task | Feature representation | No. subjects | No. observations | No. predictors |
| --- | --- | --- | --- | --- | --- |
| Alcohol initiation | Binary | Full | 2,620 | 7,860 | 786 |
| Alcohol initiation | Binary | Cluster | 2,620 | 7,860 | 686 |
| Alcohol initiation | Binary | PCA | 2,620 | 7,860 | 655 |
| Alcohol initiation | Binary | Weighted | 2,620 | 7,860 | 655 |
| Any substance initiation | Binary | Full | 2,620 | 7,860 | 786 |
| Any substance initiation | Binary | Cluster | 2,620 | 7,860 | 686 |
| Any substance initiation | Binary | PCA | 2,620 | 7,860 | 655 |
| Any substance initiation | Binary | Weighted | 2,620 | 7,860 | 655 |
| Cannabis initiation | Binary | Full | 2,620 | 7,860 | 261 |
| Cannabis initiation | Binary | Cluster | 2,620 | 7,860 | 211 |
| Cannabis initiation | Binary | PCA | 2,620 | 7,860 | 185 |
| Cannabis initiation | Binary | Weighted | 2,620 | 7,860 | 191 |
| Externalizing behavior | Continuous | Full | 2,621 | 11,108 | 122 |
| Externalizing behavior | Continuous | Cluster | 2,621 | 11,108 | 90 |
| Externalizing behavior | Continuous | PCA | 2,621 | 11,108 | 62 |
| Externalizing behavior | Continuous | Weighted | 2,621 | 11,108 | 78 |
| Internalizing behavior | Continuous | Full | 2,621 | 11,108 | 87 |
| Internalizing behavior | Continuous | Cluster | 2,621 | 11,108 | 71 |
| Internalizing behavior | Continuous | PCA | 2,621 | 11,108 | 62 |
| Internalizing behavior | Continuous | Weighted | 2,621 | 11,108 | 66 |
| Nicotine initiation | Binary | Full | 2,620 | 7,860 | 274 |
| Nicotine initiation | Binary | Cluster | 2,620 | 7,860 | 196 |
| Nicotine initiation | Binary | PCA | 2,620 | 7,860 | 153 |
| Nicotine initiation | Binary | Weighted | 2,620 | 7,860 | 167 |

**Table 2.**
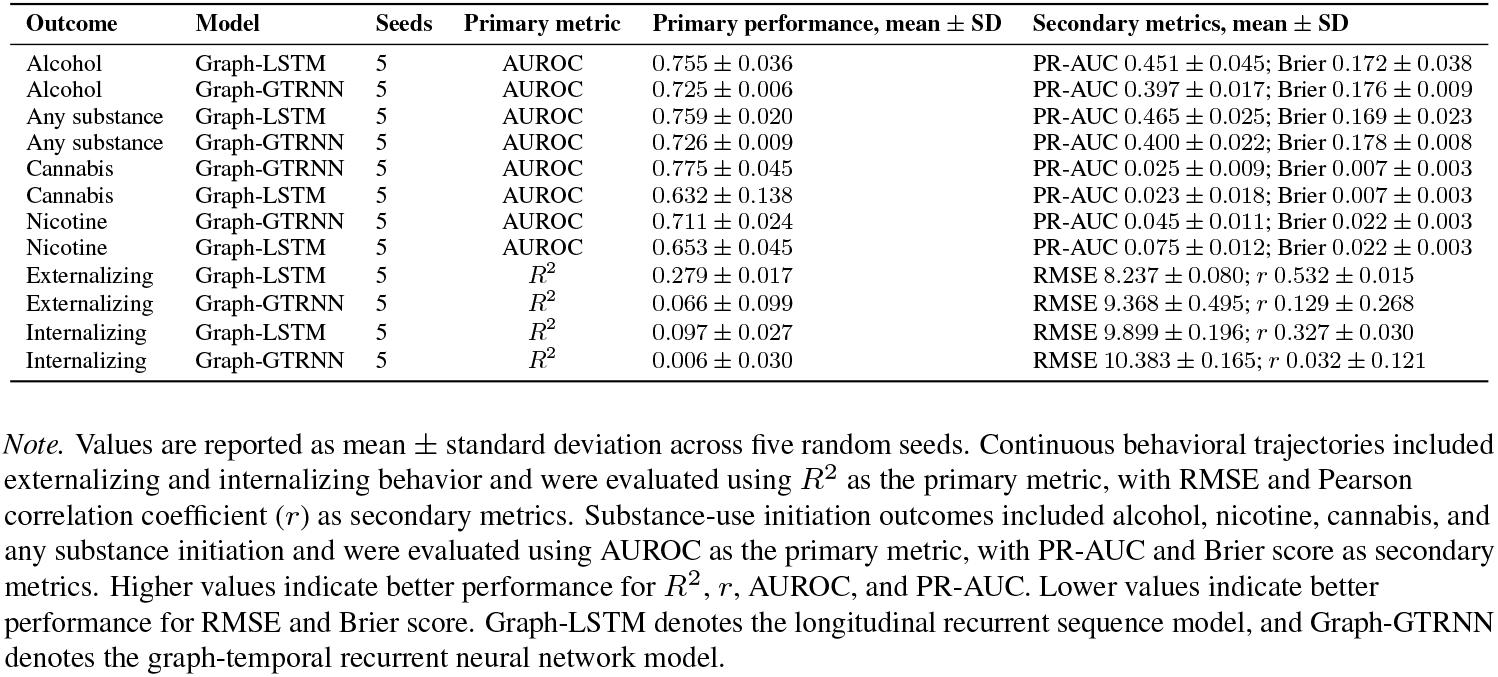
Predictive performance of Graph-LSTM and Graph-GTRNN models across continuous behavioral trajectories and substance-use initiation outcomes.

| Outcome | Model | Seeds | Primary metric | Primary performance, mean ± SD | Secondary metrics, mean ± SD |
| --- | --- | --- | --- | --- | --- |
| Alcohol | Graph-LSTM | 5 | AUROC | 0.755 ± 0.036 | PR-AUC 0.451 ± 0.045; Brier 0.172 ± 0.038 |
| Alcohol | Graph-GTRNN | 5 | AUROC | 0.725 ± 0.006 | PR-AUC 0.397 ± 0.017; Brier 0.176 ± 0.009 |
| Any substance | Graph-LSTM | 5 | AUROC | 0.759 ± 0.020 | PR-AUC 0.465 ± 0.025; Brier 0.169 ± 0.023 |
| Any substance | Graph-GTRNN | 5 | AUROC | 0.726 ± 0.009 | PR-AUC 0.400 ± 0.022; Brier 0.178 ± 0.008 |
| Cannabis | Graph-GTRNN | 5 | AUROC | 0.775 ± 0.045 | PR-AUC 0.025 ± 0.009; Brier 0.007 ± 0.003 |
| Cannabis | Graph-LSTM | 5 | AUROC | 0.632 ± 0.138 | PR-AUC 0.023 ± 0.018; Brier 0.007 ± 0.003 |
| Nicotine | Graph-GTRNN | 5 | AUROC | 0.711 ± 0.024 | PR-AUC 0.045 ± 0.011; Brier 0.022 ± 0.003 |
| Nicotine | Graph-LSTM | 5 | AUROC | 0.653 ± 0.045 | PR-AUC 0.075 ± 0.012; Brier 0.022 ± 0.003 |
| Externalizing | Graph-LSTM | 5 | $R^2$ | 0.279 ± 0.017 | RMSE 8.237 ± 0.080; $r$ 0.532 ± 0.015 |
| Externalizing | Graph-GTRNN | 5 | $R^2$ | 0.066 ± 0.099 | RMSE 9.368 ± 0.495; $r$ 0.129 ± 0.268 |
| Internalizing | Graph-LSTM | 5 | $R^2$ | 0.097 ± 0.027 | RMSE 9.899 ± 0.196; $r$ 0.327 ± 0.030 |
| Internalizing | Graph-GTRNN | 5 | $R^2$ | 0.006 ± 0.030 | RMSE 10.383 ± 0.165; $r$ 0.032 ± 0.121 |

*Note.* Values are reported as mean ± standard deviation across five random seeds. Continuous behavioral trajectories included externalizing and internalizing behavior and were evaluated using $R^2$ as the primary metric, with RMSE and Pearson correlation coefficient ( $r$ ) as secondary metrics. Substance-use initiation outcomes included alcohol, nicotine, cannabis, and any substance initiation and were evaluated using AUROC as the primary metric, with PR-AUC and Brier score as secondary metrics. Higher values indicate better performance for $R^2$ , $r$ , AUROC, and PR-AUC. Lower values indicate better performance for RMSE and Brier score. Graph-LSTM denotes the longitudinal recurrent sequence model, and Graph-GTRNN denotes the graph-temporal recurrent neural network model.

**Table 3.** Group differences in graph-derived latent energy across behavioral and substance-use outcomes.

| Outcome | Outcome type | Comparison | *n* risk/event | *n* comparison | Mean energy | Final energy | Energy change | Main pattern |
| --- | --- | --- | --- | --- | --- | --- | --- | --- |
| Externalizing | Continuous | High-symptom vs lower-symptom | 143 ± 5 | 382 ± 5 | +0.016 | +0.054 | +0.019 | Clear higher energy in high-symptom group |
| Internalizing | Continuous | High-symptom vs lower-symptom | 137 ± 5 | 388 ± 5 | +0.012 | +0.042 | +0.014 | Clear higher energy in high-symptom group |
| Alcohol initiation | Binary initiation | Event/case vs no-event | 203 ± 9 | 321 ± 9 | +0.002 | +0.004 | +0.003 | Small event-related increase |
| Any substance initiation | Binary initiation | Event/case vs no-event | 214 ± 9 | 310 ± 9 | +0.000 | +0.001 | +0.001 | Minimal group separation |
| Cannabis initiation | Binary initiation | Event/case vs no-event | 9 ± 3 | 515 ± 3 | +0.003 | +0.006 | +0.003 | Small but unstable difference |
| Nicotine initiation | Binary initiation | Event/case vs no-event | 23 ± 3 | 501 ± 3 | -0.000 | +0.003 | +0.003 | Mixed/weak difference |

*Note.* Values are reported as mean ± standard deviation across five random seeds. For continuous behavioral outcomes, the high-risk group was defined as the high-symptom group, and the comparison group was defined as the lower-symptom group. For substance-use initiation outcomes, the high-risk group was defined as the event/case group, and the comparison group was defined as the no-event group. $\Delta$ values represent the difference between the high-risk/event group and the comparison group. Positive $\Delta$ values indicate higher graph-derived latent energy in the high-risk/event group. Mean energy summarizes average latent graph-state energy over time, final energy summarizes latent energy at the final observed time point, and energy change summarizes temporal accumulation or change in latent energy. Continuous outcomes show clear and consistent separation: high-symptom groups have higher mean energy, higher final energy, and greater energy increase. Binary outcomes show weaker and less stable separation: event/case groups are only slightly different from no-event groups. For nicotine, mean energy is essentially the same or slightly lower, but final energy and energy-change differences are slightly higher. Because cannabis and nicotine have very small event groups, these results should be interpreted cautiously.

Internalizing behavior was predicted with lower accuracy than externalizing behavior. The best internalizing model was again Graph-LSTM, which achieved a mean *R*^2^ of 0.097 ± 0.027, RMSE of 9.899 ± 0.196, and correlation of 0.327 ± 0.030. Graph-GTRNN showed limited predictive performance for internalizing behavior, with a mean *R*^2^ of 0.006 ± 0.030 and correlation of 0.032 ± 0.121. These results suggest that the longitudinal recurrent sequence model captured meaningful variation in continuous behavioral trajectories, particularly for externalizing behavior, while internalizing behavior remained more difficult to predict.

### 3.3 Predictive performance across substance-use initiation outcomes

For substance-use initiation outcomes, model performance varied by phenotype (Table 2). Alcohol initiation and any substance initiation were best predicted by Graph-LSTM. For alcohol initiation, Graph-LSTM achieved a mean AUROC of 0.755±0.036, PR-AUC of 0.451±0.045, and Brier score of 0.172±0.038. Graph-GTRNN also performed above chance for alcohol initiation but showed lower discrimination, with a mean AUROC of 0.725 ± 0.006 and PR-AUC of 0.397 ± 0.017.

For any substance initiation, Graph-LSTM achieved a mean AUROC of 0.759 ± 0.020, PR-AUC of 0.465 ± 0.025, and Brier score of 0.169 ± 0.023. Graph-GTRNN achieved a lower mean AUROC of 0.726 ± 0.009 and PR-AUC of 0.400 ± 0.022. These findings indicate that for more prevalent initiation outcomes, the LSTM-based temporal model provided the strongest predictive performance.

In contrast, Graph-GTRNN performed better for cannabis and nicotine initiation. For cannabis initiation, Graph-GTRNN achieved the highest AUROC among all initiation outcomes, with a mean AUROC of 0.775 ± 0.045. Graph-LSTM showed lower and more variable performance for cannabis initiation, with a mean AUROC of 0.632 ± 0.138. For nicotine initiation, Graph-GTRNN also outperformed Graph-LSTM, achieving a mean AUROC of 0.711 ± 0.024 compared with 0.653 ± 0.045 for Graph-LSTM.

However, cannabis and nicotine initiation had very low event rates, approximately 0.7% and 2.3%, respectively. Therefore, although AUROC values were moderate to high, PR-AUC values remained low for these rare outcomes. This suggests that the models were able to rank subjects by relative risk, but rare-event prediction remained challenging in terms of precision-recall performance.

### 3.4 Best-performing model by outcome

Across the six outcomes, Graph-LSTM was the best-performing model for externalizing behavior, internalizing behavior, alcohol initiation, and any substance initiation. Graph-GTRNN was the best-performing model for cannabis and nicotine initiation. This pattern indicates that the simpler sequence-based recurrent model was most effective for dense continuous trajectories and more common initiation outcomes, whereas the graph-recurrent model may provide advantages for sparse or rare event outcomes where relational system structure may help separate high-risk individuals.

### 3.5 Graph-state and latent energy summaries

To examine whether DynoSys 2.0 captured interpretable latent system states, we summarized graph-derived energy measures from Graph-GTRNN. For continuous behavioral outcomes, high-symptom groups showed consistently higher energy than lower-symptom groups. In externalizing behavior, the high-symptom group had higher mean energy than the lower-symptom group, 0.082 versus 0.067, and higher final energy, 0.161 versus 0.106. The high-symptom externalizing group also showed greater positive energy change over time, with mean energy delta of 0.018 compared with approximately 0.000 in the lower-symptom group.

A similar pattern was observed for internalizing behavior. The high-symptom internalizing group showed higher mean energy than the lower-symptom group, 0.076 versus 0.064, and higher final energy, 0.152 versus 0.110. The high-symptom group also showed greater energy accumulation over time, with mean energy delta of 0.016 compared with 0.002 in the lower-symptom group. These results suggest that higher symptom burden is associated with elevated latent system energy and greater dynamic instability in the graph-recurrent representation.

For substance-use initiation outcomes, energy differences between event/case and no-event subjects were smaller. Alcohol, cannabis, and any substance initiation showed slightly higher mean or final energy among event/case subjects than no-event subjects, whereas nicotine showed minimal group separation. These results suggest that graph-state summaries may be more interpretable for continuous symptom trajectories than for rare initiation outcomes, although modest event-related energy differences were observed for some substance-use phenotypes.

Overall, the graph-state analyses provide complementary evidence that DynoSys 2.0 captures latent dynamic system properties beyond standard predictive performance. In particular, elevated energy among high-symptom externalizing and internalizing groups supports the interpretation of behavioral symptoms as emergent states of a dynamic, multi-domain system.

## 4 Discussion

In this study, we developed and evaluated DynoSys 2.0, a longitudinal dynamic systems framework for modeling behavioral development using multi-domain inputs. The goal of this framework was to represent human behavior as the result of interactions among biological, environmental, developmental, and system-level factors over time. Across six behavioral outcomes and five random seeds, DynoSys 2.0 generated stable longitudinal sequence inputs and allowed us to compare two temporal modeling approaches: a sequence-based recurrent model, Graph-LSTM, and a graph-recurrent model, Graph-GTRNN.

A major finding was that model performance depended strongly on the type of outcome. For continuous behavioral outcomes, externalizing behavior was predicted more accurately than internalizing behavior. The best externalizing model achieved a mean *R*^2^ of 0.279 and a correlation of 0.532 across random seeds. In contrast, the best internalizing model achieved a lower mean *R*^2^ of 0.097 and a correlation of 0.327. This suggests that the current available longitudinal predictors may capture externalizing-related developmental processes better than internalizing-related processes. Externalizing behaviors may be more visible and more strongly linked to measured environmental and developmental features, while internalizing symptoms may be more heterogeneous, less directly observable, and influenced by factors that were not fully measured in the current data.

For substance-use initiation outcomes, performance also varied across phenotypes. Alcohol initiation and any substance initiation were best predicted by the LSTM model, with AUROC values of approximately 0.76. These results suggest that longitudinal developmental patterns contain useful information for predicting later substance-use initiation. In contrast, cannabis and nicotine initiation were best predicted by the graph-recurrent model. This may indicate that graph-based modeling provides additional value for sparse or rare-event outcomes, where the number of positive cases is limited.

However, the graph-recurrent model did not outperform the LSTM model for all outcomes. Instead, the results suggest a more balanced interpretation. The simpler LSTM model appeared to be sufficient, and sometimes better, for continuous outcomes and more common initiation outcomes. The graph-recurrent model may be more useful for rare initiation outcomes such as cannabis and nicotine. Therefore, DynoSys 2.0 should not be interpreted as a framework where one model is always superior. Rather, it should be viewed as a flexible dynamic systems framework in which different temporal architectures may work better for different types of behavioral outcomes.

In addition to prediction, DynoSys 2.0 also provided graph-derived summaries of latent system dynamics. Although detailed node-to-node connectivity differences were not the main focus of the current study, graph-state and energy analyses showed outcome-specific differences in latent system states. These differences were clearest for continuous behavioral symptoms, especially externalizing and internalizing behavior. High-symptom individuals showed higher mean energy, higher final energy, and greater accumulated energy over time compared with lower-symptom individuals.

These graph-state findings support the main systems-level hypothesis of the study. If genes and environmental exposures are viewed as system inputs, and behavior is viewed as the system output, then the latent recurrent graph state can be interpreted as an intermediate system representation. The finding that high-symptom individuals showed elevated latent energy suggests that behavioral symptoms may reflect more activated, unstable, or dysregulated system states over development. This interpretation was strongest for externalizing and internalizing behavior, where high- and low-symptom groups showed clear separation in graph-derived energy summaries.

For initiation outcomes, graph-state differences between event and no-event groups were weaker. This may be due to the low prevalence of several initiation outcomes, especially cannabis and nicotine, and also because initiation outcomes are discrete events rather than continuous symptom trajectories. Rare-event prediction is difficult because a model can show a moderate or high AUROC while still having a low PR-AUC. In this case, AUROC suggests that the model can rank individuals by risk better than chance, but low PR-AUC means that accurately identifying future positive cases remains challenging. Therefore, the initiation results should be interpreted carefully.

Together, these results provide partial support for the DynoSys 2.0 hypothesis. The findings show that longitudinal multi-domain information can predict both behavioral trajectories and substance-use initiation. They also show that model-derived graph states can distinguish high-symptom from lower-symptom individuals. These results are consistent with the view that behavior is not determined by one isolated factor. Instead, behavioral outcomes may emerge from dynamic interactions among biological susceptibility, environmental exposure, developmental timing, and latent system-state changes.

At the same time, the results also show the need for caution. Predictive performance was stronger for externalizing behavior than for internalizing behavior, suggesting that the current feature set may better capture externalizing-related risk processes. Rare-event outcomes such as cannabis and nicotine initiation had low PR-AUC despite moderate to high AUROC, showing that class imbalance remains a major challenge. In addition, graph energy measures are model-derived latent summaries, not direct biological measurements. They should therefore be interpreted as computational indicators of system dynamics, rather than direct physiological energy states. Finally, because this study used observational data, the predictive patterns and graph-state differences should not be interpreted as causal effects without further causal modeling or experimental validation.

Overall, DynoSys 2.0 provides a systems-level approach for studying behavioral development across both continuous and event-based outcomes. The framework supports three main conclusions. First, longitudinal multi-domain data contain meaningful signal for predicting behavioral trajectories and substance-use initiation. Second, the best temporal model depends on the outcome type: LSTM models performed best for dense continuous outcomes and more common initiation outcomes, while graph-recurrent models showed advantages for rare initiation outcomes. Third, graph-derived latent energy states distinguished high-symptom from lower-symptom individuals, supporting the view that behavioral symptoms can be represented as emergent states of a dynamic developmental system.

### 4.1 Limitations

Several limitations should be considered. First, graph construction depends on modeling assumptions. These include how features are mapped to nodes and how relationships among domains are defined. Different graph definitions may lead to different latent state summaries.

Second, the data are observational. Therefore, the associations identified by the models should not be interpreted as causal effects. Additional causal modeling or experimental validation would be needed to support causal claims.

Third, graph energy measures are latent model-derived quantities. They are not direct biological measurements. Therefore, they should be interpreted as computational summaries of system activity, rather than as physical or physiological energy states.

Fourth, rare-event outcomes remain difficult to model. Cannabis and nicotine initiation had low event rates, which reduced precision-recall performance and made group-level graph-state comparisons less stable. Although the models could rank risk above chance, accurate identification of rare positive cases remains challenging.

Fifth, the current study showed clear graph-state differences for continuous symptoms, but it did not fully establish detailed graph connectivity patterns, stable versus unstable nodes, or specific intervention targets. Future analyses should examine node-level and edge-level graph dynamics in more detail before making stronger claims about mechanisms or intervention points.

## 5 Conclusion

In conclusion, DynoSys 2.0 provides a longitudinal dynamic systems framework for modeling behavioral development using multi-domain inputs. The framework achieved meaningful prediction across continuous behavioral trajectories and substance-use initiation outcomes. It also provided interpretable graph-state summaries of latent system dynamics.

The strongest evidence for the systems-level interpretation came from externalizing and internalizing behavior. In these outcomes, high-symptom individuals showed elevated latent energy and greater energy accumulation over time compared with lower-symptom individuals. These findings support the idea that behavioral risk reflects dynamic interactions among system components, rather than isolated predictors alone.

Overall, DynoSys 2.0 suggests that behavioral outcomes can be modeled as emergent states of a developmental system shaped by biological, environmental, and temporal processes. Future work should extend this framework to larger samples, improve rare-event prediction, include additional neurobiological features, and test whether latent system-state measures can help identify modifiable intervention targets.

## 7 Author Contributions

Mengman Wei conceived the study, designed the analytical framework, performed all data processing, statistical analyses, and computational modeling, and drafted the manuscript. All code implementation, data curation, and result interpretation were conducted by Mengman Wei.

Qian Peng provided supervision, general guidance, resource support, and funding acquisition.

## 8 Preprint Notice

This manuscript is a preprint and has not yet undergone peer review. The content is shared to disseminate findings and establish precedence. Additional analyses and revisions may be incorporated in future versions.

## 9 Data availability

### Code

The analysis code and scripts used in this study are freely available at the following GitHub repository: https://github.com/mw742/DynoSys2.0.

### Data

This study uses data from the Adolescent Brain Cognitive Development (ABCD) Study (https://abcdstudy.org), held in the NIMH Data Archive (NDA). The ABCD data release used was version 5.1. The study is supported by the National Institutes of Health (NIH) and additional federal partners under multiple award numbers, including U01DA041048 and U01DA050987. The full list of funders is available at https://abcdstudy.org/federal-partners.html.

